# Uncovering the locus coeruleus: comparison of localization methods for functional analysis

**DOI:** 10.1101/2020.05.12.092320

**Authors:** Verónica Mäki-Marttunen, Thomas Espeseth

## Abstract

Functional neuroimaging of small brainstem structures in humans is gaining increasing interest due to their potential importance in aging and many clinical conditions. Researchers have used different methods to measure activity in the locus coeruleus (LC), the main noradrenergic nucleus in the brain. However, the reliability of the different methods for identifying this small structure is unclear. In the present article, we compared four different approaches to estimate localization of the LC in a large sample (N = 98): 1) a probabilistic map from a previous study, 2) masks segmented from neuromelanin-sensitive scans, 3) components from a masked-independent components analysis of the functional data, and 4) a mask from pupil regression of the functional data. The four methods have been used in the community and find some support as reliable ways of assessing the localization of LC *in vivo* in humans by using functional imaging. We report several measures of similarity between the LC masks obtained from the different methods. In addition, we compare the similarity between functional connectivity maps obtained from the different masks. We conclude that sample-specific masks appear more suitable than masks from a different sample, that masks based on structural versus functional methods may capture different portions of LC, and that, at the group level, the creation of a “consensus” mask using more than one approach may give a better estimate of LC localization.

## 1. Introduction

Advances in imaging research have opened opportunities to study small brainstem structures in humans (Beissner et al., 2014; Sclocco et al., 2018). Between them, the locus coeruleus (LC) has gained increasing interest in the past decade. LC projects to most of the cortical and subcortical areas (Szabadi, 2013). A wealth of findings in animal studies suggest that the LC-norepinephrine (LC-NE) system participates in the coordination of brain processing, enabling the expression of states such as cognition, attention and quiet wake (Berridge & Waterhouse, 2003). This has direct implications for human neuroscience. In addition, the LC-NE system is increasingly related to brain pathologies, such as Alzheimer’s disease (Friedman et al., 1999; Heneka et al., 2010; Weinshenker, 2008, 2018), Parkinson (Bari et al., 2020; Gesi et al., 2000), and dementia (Jacobs et al., 2015; Mather & Harley, 2016). These clinical conditions present neuronal loss in LC and LC hyperactivity as early markers, and thus the ability to measure LC structural and functional integrity has great potential for early diagnosis of neurodegenerative pathologies (Betts et al., 2019). A great obstacle in deepening the study of the LC in humans is its small size, which makes it difficult to localize with standard imaging methods.

The LC nucleus has steadily gained increasing interest in human research. New theories and revision/extension of older theories about its involvement in attention, learning and cognition have been proposed in recent years (Aston-Jones & Waterhouse, 2016; Guedj et al., 2017; Sara & Bouret, 2012), accompanied by experimental work using fMRI. A number of studies using MRI have tested LC integrity in conditions characterized by cognitive deficits, which emphasizes the clinical relevance of research on LC. Even though the limitations of early methodological approaches have been identified and improved in recent work (Astafiev et al., 2010), there is still a need for a thorough assessment and comparison of available methods.

Two main goals may be identified as to obtain a reliable localizer of LC in functional studies: one is at the group level, where a reliable LC mask allows to determine whether significant activity in LC is observed in a group statistical map. The other goal is to use LC masks to extract activity (either task-related or for functional connectivity analysis) within each individual’s LC. While the two are related, they may be seen as two extremes in a continuum (**Figure 1**): while the former requires a constrained “consensus” region where to reliably determine presence of true LC activity, ignoring “borders” or regions with higher spatial variability, the latter emphasizes individual specificity of LC localization.

**Figure 1.**
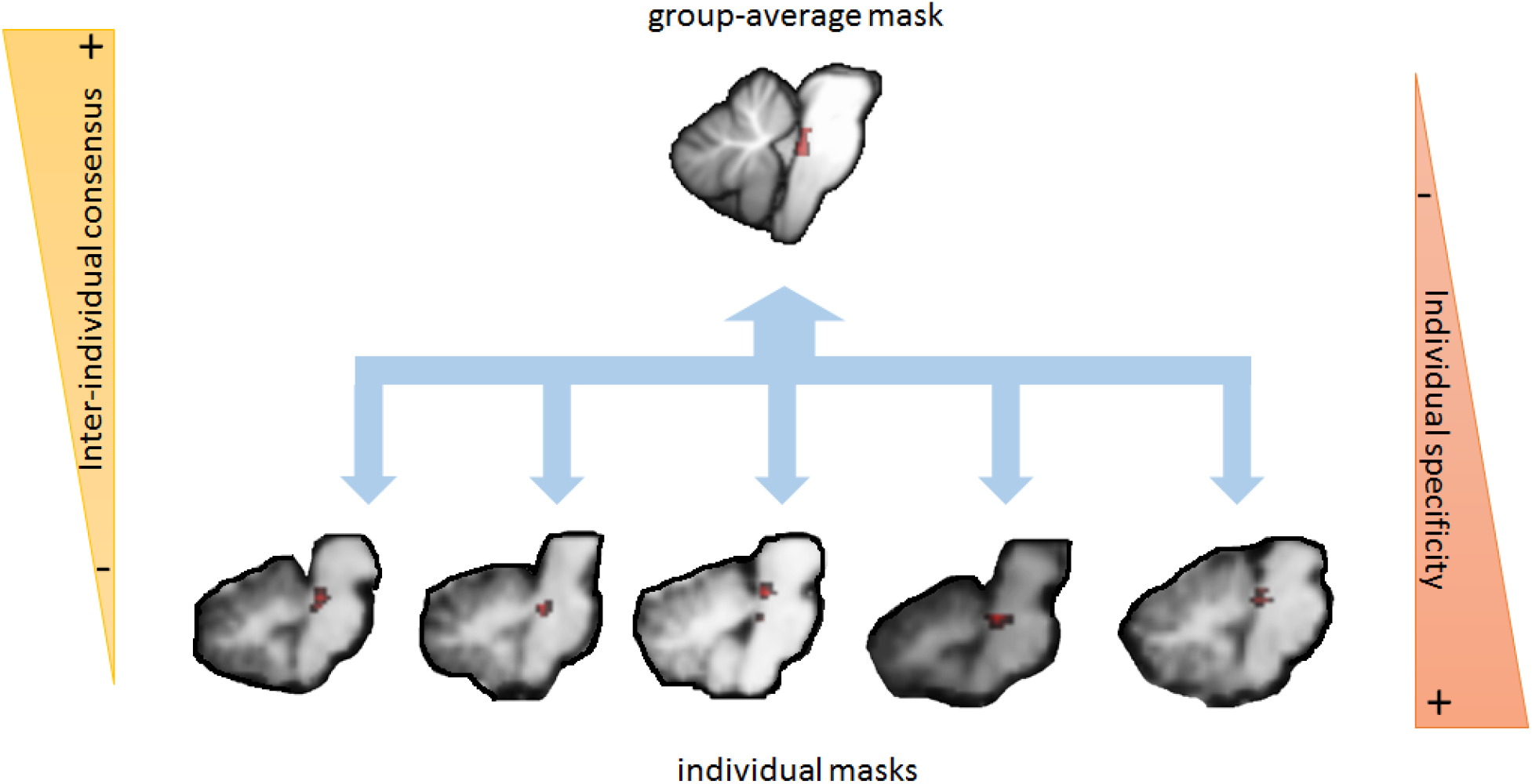
Illustration of approaches for LC localization. Group approaches emphasize voxels of highest probability of containing LC at the group level. Individual approaches focus on identifying LC-containing voxels in the individual brainstem.

A recent systematic review of imaging studies of LC (Liu et al., 2017) highlights the large variety in the methods used to identify the region that more closely corresponds to the LC. A majority of the reviewed studies used coordinates or masks from previous studies, which may compromise the degree to which the MRI signal had a true LC origin in the study-specific samples. The peak coordinates reported for the LC in functional studies reveal great variability and cover a larger region than the one where LC could be truly located (Liu et al., 2017). A point raised by Liu et al. was to what extent the variability in LC position across individuals can influence variability in LC localization across studies, and to what extent this could be due to methodological issues.

Studies aiming to characterize LC at the individual level have used a special structural sequence in which tissues that contain high density of the molecule neuromelanin (as is the case of LC and substantia nigra) appear as high intensity regions in the images (Keren et al., 2009; Nakamura & Sugaya, 2014; Sasaki et al., 2008). Comparison of MRI and post-mortem histology confirmed that high intensity voxels using neuromelanin-sensitive sequences correspond to regions with high concentration of neuromelanin and tyrosine hydroxylase positive cells, both of which are found in the LC (Keren et al., 2015). This is possibly the best way of assessing the position of LC in individual brains in vivo, although some variation in the methodology is also present. Recent work suggests that the manual identification of LC in the neuromelanin sequence has moderate to good reliability (Langley et al., 2017; Tona et al., 2017). Most of the studies using the neuromelanin sequence were purely structural (i.e. measured peak intensity of neuromelanin scan). A few functional studies used this method for individualized LC localization to test hypotheses regarding LC activity during different cognitive tasks (Clewett et al., 2018; Krebs et al., 2018), including our own work (Mäki-Marttunen et al., 2019b; Mäki-Marttunen et al., 2020). Recent studies have aimed for improved manual delineation of LC in neuromelanin scans by using a semi-automatic segmentation of LC (Betts et al., 2017; Dahl et al., 2019). In this approach, the LC is drawn on a group average image of aligned neuromelanin scans, and this mask is used as a search space for high-intensity voxels in the individual scans.

In addition to reported data and the neuromelanin sequence, two other methods can be found in the literature that in principle allow to identify the position of LC. Both are based on functional analyses. The first method involves an independent component analysis (ICA) of individual functional data (i.e. resting state) limited to the brainstem, and identifies the independent component that overlaps with the expected position of LC. This method, called masked ICA (mICA), has been proposed for studies on the activity in brainstem nuclei (Beissner et al., 2014). The second method is based on the finding in animal studies that pupil size, after removing luminance effects, co-varies with activity in LC (Joshi et al., 2016). In this way, regressing the pupil size time course with BOLD activity would give a significant spot in the LC (Murphy et al., 2014).

In sum, several methods have been used to more or less directly assess LC localization in humans. However, the validity of each of them is currently unclear. This is an important issue to settle before more imaging research focuses on this nucleus. In the present study, we compared four methods to localize LC: 1) a probabilistic mask from a previous study (Keren et al., 2009); 2) a neuromelanin-sensitive sequence that allows to detect the LC as a zone of high intensity, both using manual and semi-automatic segmentations; 3) a masked independent components analysis on functional data; 4) a mask based on pupil size time-series correlation. We compared the characteristics of the obtained masks: size, position, and signal correlation. We also aimed to replicate functional connectivity (FC) patterns of LC from previous studies using the different methods of LC identification. We compared the functional connectivity maps resulting from the different LC masks. We argue that if the masks present overlap and the obtained connectivity maps are comparable, any of the methods would in principle reliably identify the LC. In the era of big data, such knowledge is needed to validate the study of small brainstem structures applied to large datasets with as little manual involvement as possible. These results will also have implications for the study of other small structures in the brainstem. The results intend to work towards a consensus about reliable LC localization and extraction of activity in humans.

## 2. Methods

### 2.1 Dataset

101 participants (age: 26 ± 5, range: 19-42, N female = 63) were recruited in the University of Oslo. The work was carried out in accordance with the Declaration of Helsinki. Informed written consent was obtained in accordance with the protocols approved by the local Ethics Committee, and all participants were given 100 NOK per hour for their participation. None of the participants had a present or past history of serious disorders. Three participants were excluded from the functional analysis for bad reconstruction of images from the scanner (1) or because the data was incomplete (2). A working memory test (letters-digits span task) was administered outside the scanner.

### 2.2 Image acquisition and preprocessing

MRI sessions took place in the 3T Philips Achieva scanner (8-channel head coil) at Rikshospitalet, Oslo. Each scanning session started with an anatomical scan. The resting state paradigm lasted 10 minutes, in which 250 volumes were collected. In addition, three functional scans with an attentional task were acquired. Whole-brain functional images were acquired using a spin-echo echo-planar (EPI) sequence sensitive to blood-oxygen-level-dependent (BOLD) magnetic susceptibility (TR = 2500 ms; Flip angle = 90º; number of slices: 42; voxel size: 3 mm^3^). A neuromelanin-sensitive scan was acquired with the objective of further improving the localization of the locus coeruleus. The slices for this scan were set in order to cover the brainstem (T1-TSE, TR: 600ms, TE: 14ms, voxel dimension: 0.4×0.49×3mm, Flip angle = 90º; number of slices: 10). This scan lasted 12 minutes. During the resting state, a cross was projected on a screen positioned at the head end of the litter. Participants viewed the screen through a mirror placed on the head coil.

The functional images of each subject were first visually inspected for anomalies and then submitted to a standard preprocessing pipeline using SPM 12 implemented on MATLAB (Mathworks). Images were first corrected for time delays and realigned using 6 parameters of movement. The data were normalized to a standard template in the MNI system (image size: 75×95×75, voxel size: 2×2×2 mm^3^), and smoothed using 3 mm FWHM Gaussian kernel. Structural and functional images were loaded in the CONN toolbox (Whitfield-Gabrieli & Nieto-Castanon, 2012). Further preprocessing steps for the connectivity analysis included band-pass filtering (0.008-0.1 Hz), linear detrending and removal of mean signal from white matter and cerebro-spinal fluid.

### 2.3 Signal intensity extraction

LC signal intensity was extracted from each individual’s neuromelanin sequence according to a standard procedure (Betts et al., 2017; Clewett et al., 2016; Liu et al., 2019; Sasaki et al., 2008; Shibata et al., 2006). First, on the axial slices where LC anatomical position was expected (Keren et al., 2009; Keren et al., 2015) we identified the voxel of highest intensity in each lateral side. Then, a region of interest (ROI) was drawn as a 3 voxels x 3 voxels cross centered on that voxel using MRIcron (Rorden & Brett, 2000) drawing tools. The mean signal intensities from the left and right LC ROIs were extracted and averaged. We then drew a reference region of 10 x 10 voxels in the pontine tegmentum (PT), centered at halfway distance from the center of the two LC ROIs. The average intensity from this region was used to calculate the contrast-to-noise ratio as follows:

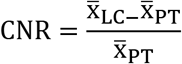

Where *x*_LC_ is the mean in LC ROIs and *x*_PT_ is the mean in the PT ROI.

### 2.4 Definition of LC masks

For the atlas-based LC (LC AT), we used the probabilistic map of LC provided by Keren et al. (2009, 2 standard deviation) as region of interest (**Figure 2**).

**Figure 2.**
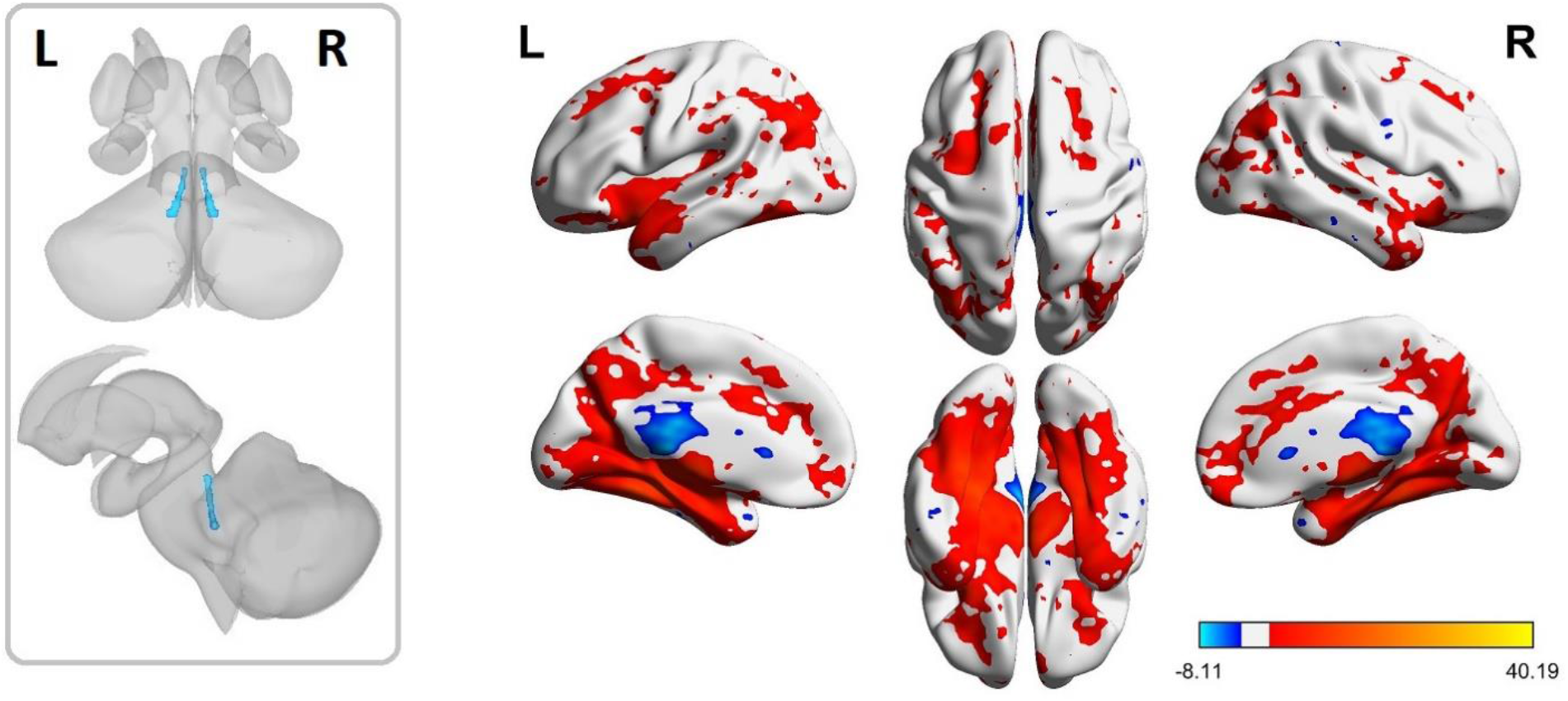
LC AT mask. Left: Probabilistic map from the study done by Keren et al. 2009. This mask (LC AT) was used as a region of interest. Right: group FC map with LC AT as seed.

For the neuromelanin-based mask (LC NM), we defined individual masks in the high-resolution T1-TSE images and used them for a region of interest analysis (**Figure 3**). The locus coeruleus of each individual was delineated on the axial slices of the neuromelanin scans. The position of the nuclei was determined in the pons as the voxels of hyperintensity on either side of the fourth ventricle. All segmentations were performed by the same experienced researcher. The T1-TSE scan of each individual was coregistered to the individual’s structural image, the structural was coregistered to the mean functional image, and the deformation field calculated during the normalization step was applied to the coregistered T1-TSE image. Correct coregistration was checked by careful visual inspection. In addition to this manual delineation of LC on the neuromelanin-sensitive sequence, we performed a semi-automatic segmentation following previously reported procedures (Betts et al., 2017; Dahl et al., 2019), adapting the code shared with us by the authors. We report the methodological procedure and the functional connectivity results of the semi-automatic procedure as Supplementary Material.

**Figure 3.**
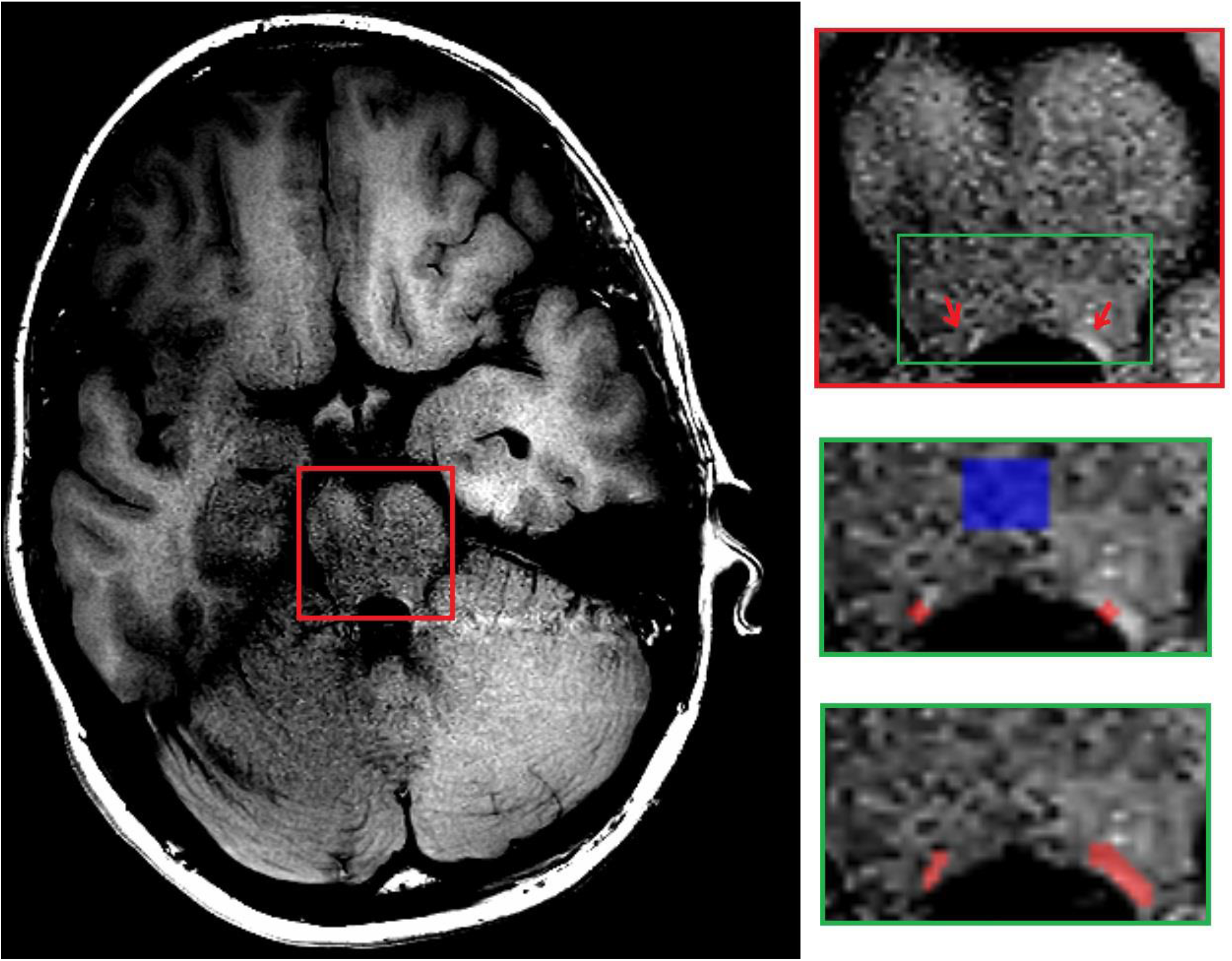
Neuromelanin-based masks definition. Example of a slice from a TSE sequence (left) on which the bilateral LC was identified as regions of high intensity in the proximity of the fourth ventricle (right, top; see full description in the main text). Contrast-to-noise ratios (CNR) were estimated based on the average intensity of a cross drawn around the voxel of maximum intensity and of a control region in the pons (right, center). Individual LC masks were drawn around the high-intensity voxels (right, bottom).

For the functional-based masks (LC mICA), we performed a masked independent component analysis (mICA) of the functional imaging data (Moher Alsady et al., 2016). Briefly, this analysis consists of a data-driven decomposition of the signal using independent component analysis (ICA), restricting the space of analysis to the brainstem and masking the rest of the brain out. This procedure allows for removing the physiological noise introduced by surrounding areas of the brainstem. Before the mICA analysis, we performed a brainstem-specific normalization. The neuroimaging data obtained after slice timing and realignment were normalized to the MNI-based cerebellum-brainstem template using the SPM toolbox SUIT 3.3 (Diedrichsen, 2006). We applied the following preprocessing steps: 1) isolation of the brainstem, 2) further manual cropping of the image, retaining only the brainstem, 3) normalization, and 4) reslice to a voxel size of 2 × 2 × 2 mm^3^. We performed a masked independent component analysis (mICA) on the functional imaging data per subject. The functional images and the cropped brainstem mask of each individual resulting from the steps using SUIT were loaded in the mICA toolbox for FSL, and the decomposition was run without setting a specific number of components. To identify the independent component or components that corresponded to LC, we calculated for each subject, the overlap of each component with the atlas mask (Keren et al., 2009), and the component or components that showed the largest overlap (more than three standard deviations) were combined in a single image. Then, the combined image of each participant was manually thresholded (this also allowed for visual inspection of the components) to leave a mask of ^~^20 voxels, and then it was binarized.

For the pupil-based mask (LC pupil), we used a subset of the data (N = 49) for which concurrent fMRI-pupillometry had been acquired during the resting state scan. For the acquisition of pupil data we used an MRI-compatible yeLink 1000 eye tracker. The pupil data was preprocessed by interpolating blinks (code adapted from Urai et al. 2017) and resampling to the number of MRI volumes. The pupil signal was used as a seed temporal course for a connectivity analysis (below). The mask of the LC was extracted from the group connectivity map of the pupil regressor after thresholding (t = 2.7).

### 2.5 LC consensus mask

A group-average LC NM mask and a group-average mICA mask were created as the average of the corresponding individual masks using imcalc function in SPM 12. Then, a threshold was applied to obtain a LC size of 120 mm^3^ (unilateral). We calculated a “consensus” LC mask by extracting the voxels that were included in at least 3 of the masks obtained from the four above-mentioned methods (AT, mean NM, mean mICA, pupil). We did this to examine the location of voxels with highest probability of belonging to the LC at the group level. We used this mask as a fifth seed in the functional connectivity analysis.

### 2.6 Functional connectivity analysis and comparison across masks

Preprocessed functional images underwent functional connectivity analysis. The average signal from the different LC masks and the pupil temporal courses were used as regressors. The Pearson correlation between the time course of each seed and the time course of each brain voxel was calculated to create individual FC maps. Then, second-level maps were obtained and thresholded at voxel-level significance of p < 0.001 and FDR corrected cluster-level significance of p < 0.05. Results were visualized using xjView toolbox (https://www.alivelearn.net/xjview) and BrainNet Viewer (Xia et al. 2013, http://www.nitrc.org/projects/bnv/). Similarity of functional connectivity maps was calculated in three different ways. First, we calculated the Pearson correlation between pairs of images; second, we created Bland-Altman plots; third, we calculated the Euler characteristic of each image and compared the curves. The code used to calculate the two last measures was adapted from Bowring et al. 2019.

## 3. Results

### 3.1 LC contrast-to-noise ratio

We first calculated the contrast-to-noise ratio (CNR) in the neuromelanin sequences to characterize LC and compare with previous studies. We found a distribution of CNR with an average of 0.099 (standard deviation: 0.034; **Figure 4**). The CNR did not correlate significantly with age (r = 0.074, p = 0.46).

**Figure 4.**
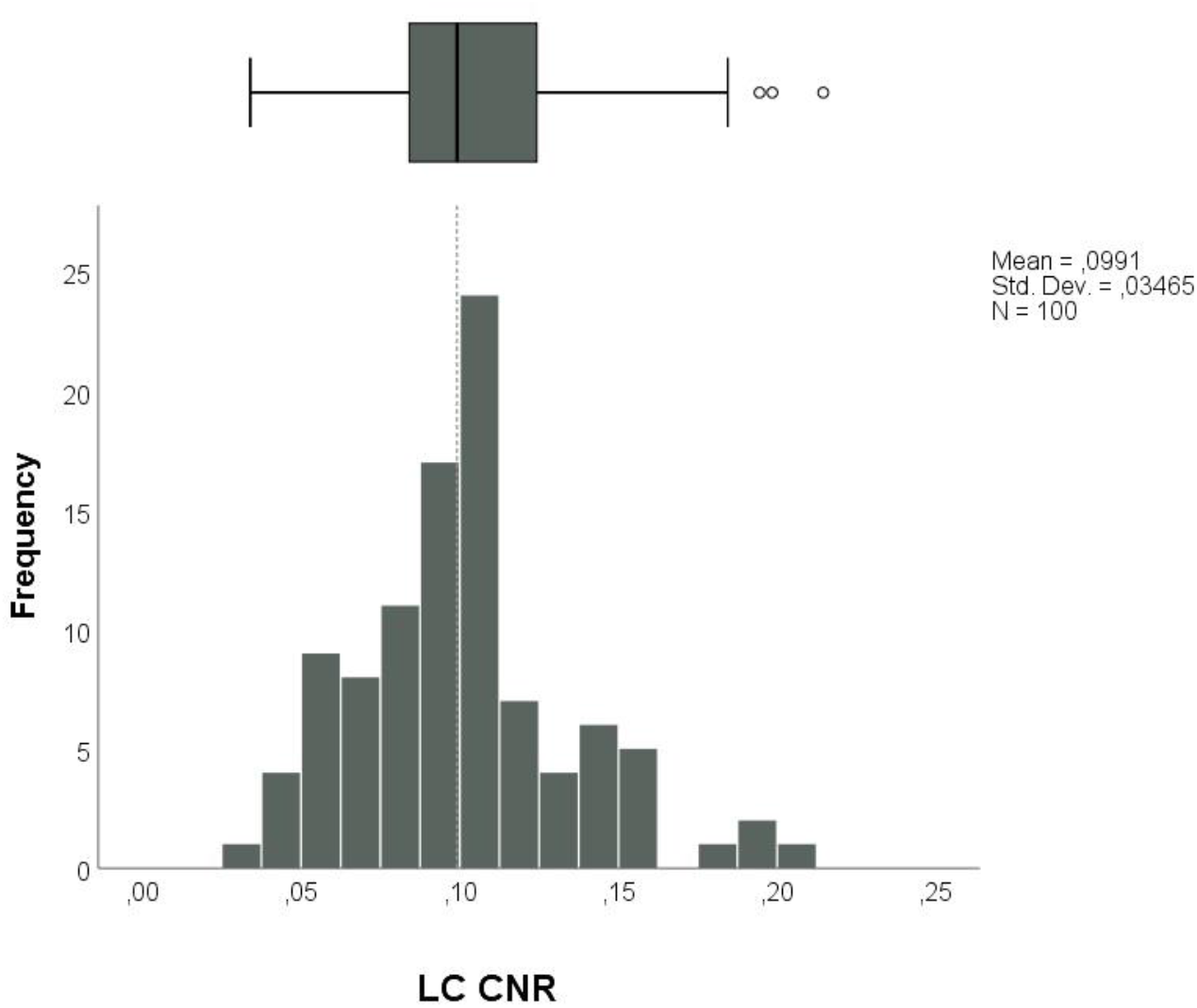
Contrast-to-noise ratio (CNR) of neuromelanin-based masks. Distribution of CNR values displayed as a histogram (bottom) and box-plot (top).

### 3.2 LC masks

The LC localization approaches analyzed in the present paper were: use of a probabilistic mask from a previous study (**LC AT**, Keren et al. 2009), individually-drawn masks on a neuromelanin-sensitive structural sequence (**LC NM**), independent components-based decomposition of brainstem signal on each individual’s data (**LC mICA**), and pupil signal regression (**LC pupil**). The LC mask from the regression of the pupil signal, was unilateral. The volume of the masks and coordinates of maximum overlap are displayed in **Table 1**. For the analyses below that involved coordinates, we used two peak coordinates of the LC AT mask that were reported to show the highest overlap at two different z values: z = −24 (AT1) and z = −27 (AT2) (Keren et al., 2009). The average volume of the individual neuromelanin-based masks (27 mm^3^ unilateral) is similar to previously reported volumes using the same method.

**Table 1.** Average volume and peak coordinate of LC masks obtained from the different methods.

| METHOD | AVERAGE VOLUME OF MASK<sup>1</sup><br>(UNILAT) MM<sup>3</sup> | PEAK COORDINATE |
| --- | --- | --- |
| LC AT | 95 | AT1: $x = -3.7, y = -37.0, z = -24^{\#}$<br>$x = 5.2, y = -36.9, z = -24^{\#}$<br>AT2: $x = -4.7, y = -37.3, z = -27^{\#}$<br>$x = 5.8, y = -37.2, z = -27^{\#}$ |
| LC NM | 27 | $x = -4, y = -38, z = -26$<br>$x = 6, y = -38, z = -26^*$ |
| LC MICA | 150 | $x = -6, y = -38, z = -27^*$<br>$x = 4, y = -38, z = -29$ |

<sup>1</sup> From literature, the expected size is between 50 and 75 mm<sup>3</sup> (unilateral; Keren et al. 2015).

|  |  |  |
| --- | --- | --- |
| LC PUPIL | 136 | $x = -6, y = -39, z = -30$ |

### 3.3 LC group-average masks

First we examined the similarity between group-average LC masks obtained from the different methods. In **Figure 5** we display the extent within the brainstem of the group-average masks, and in **Figure 6** we display the location of the peak coordinates, as in **Table 1**. Visual inspection showed an overall concordance in LC localization between methods and with previously published results, in particular on the middle portion of the nucleus. Still, some variability across methods seemed to be present, so we next analyzed the overlap between the average masks and distance between peak coordinates.

**Figure 5.**
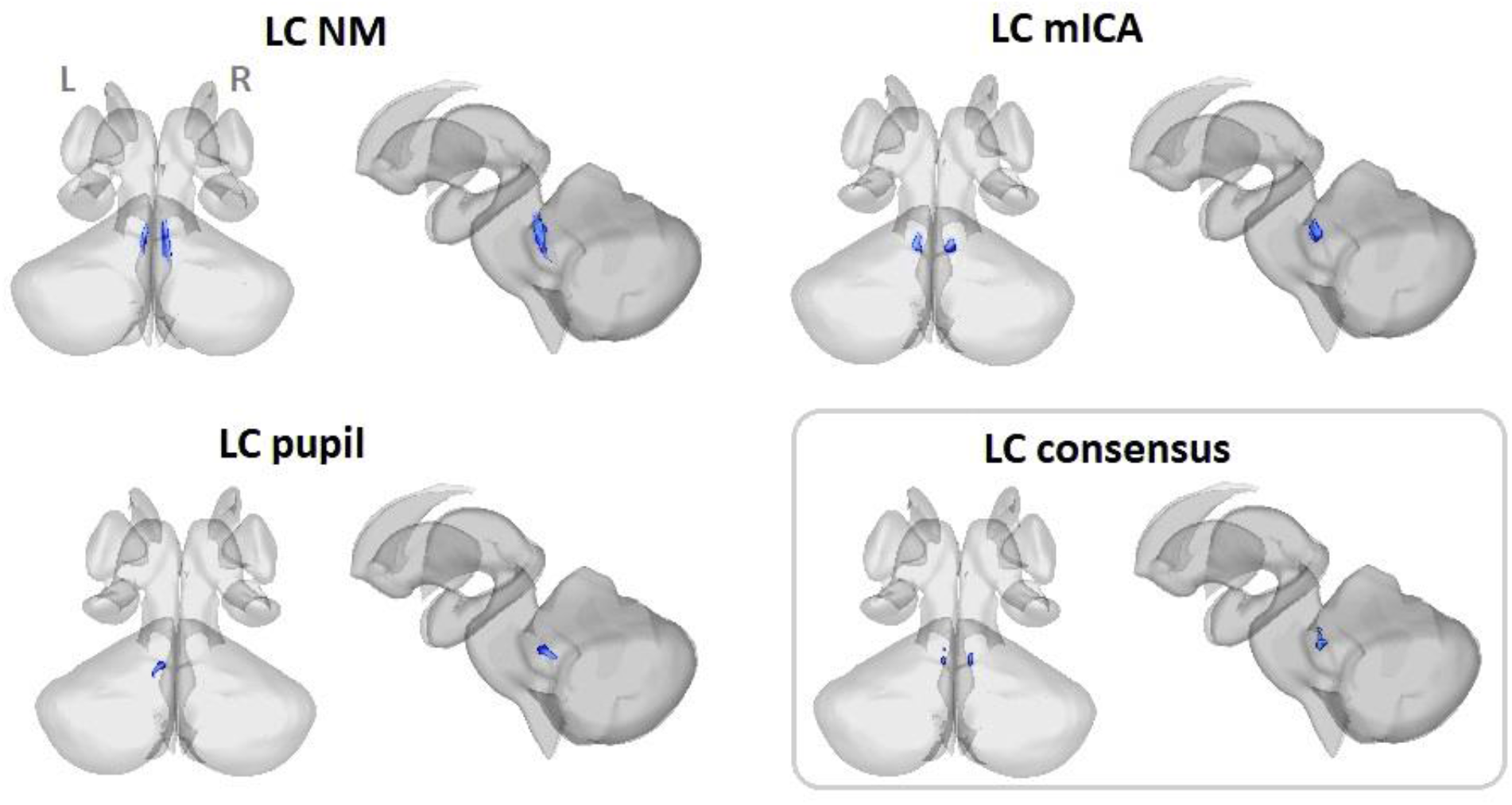
LC group-average masks obtained from the different methods. LC NM: group average mask from neuromelanin-sensitive sequences; LC mICA: group average mask from masked ICA; LC pupil: statistical map from regression of pupil covariate. LC consensus: voxels with overlap of at least three of the group LC masks from the different approaches: LC AT, LC NM, LC mICA, and LC pupil.

**Figure 6.**
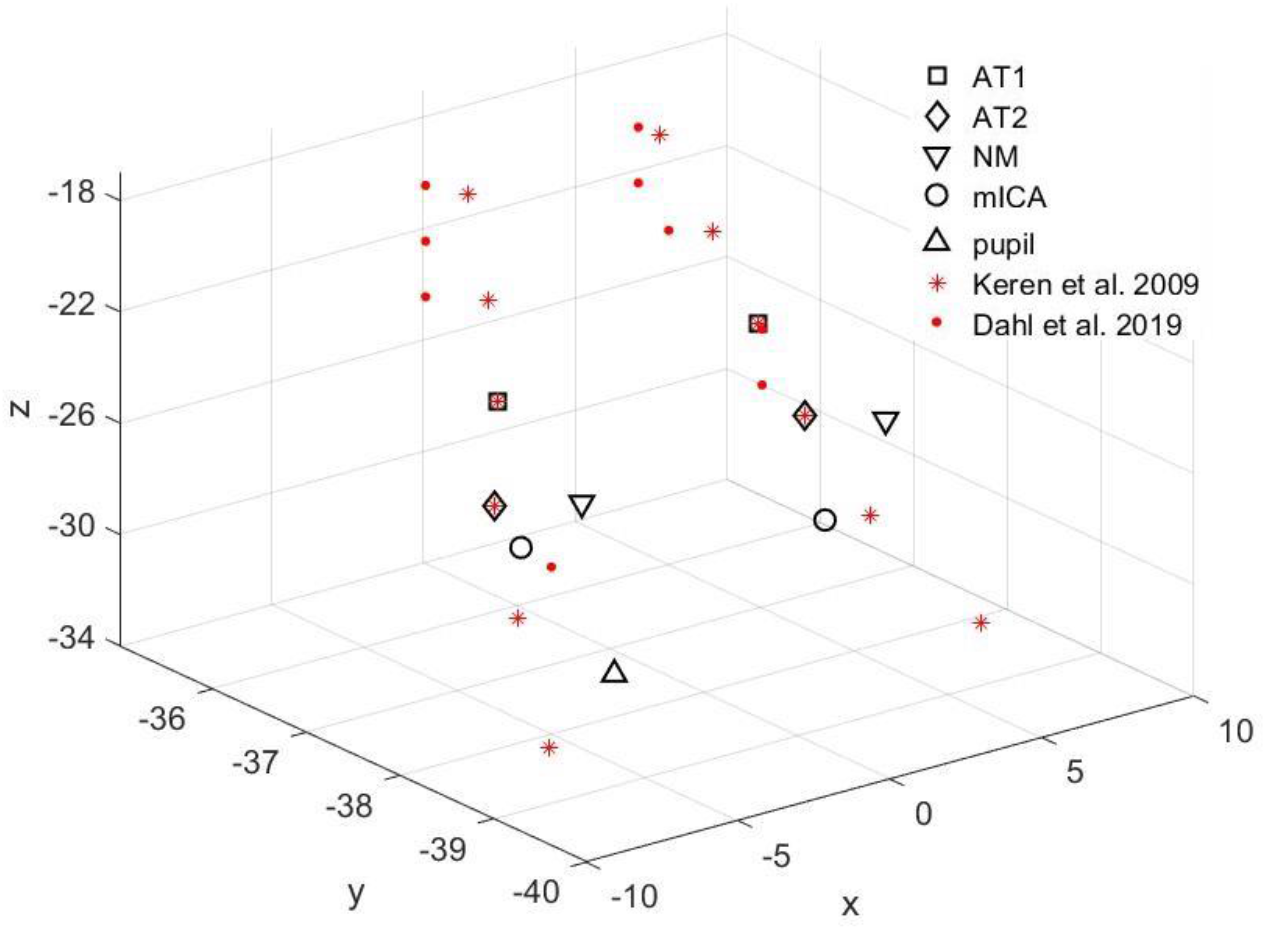
Peak coordinates of the different group-average masks (black markers). Red markers indicate maximum peak coordinates per slice in the z axis as reported in previous studies (Dahl et al., 2019; Keren et al., 2009).

The masks LC AT, LC pupil, and the group average masks of LC NM and LC mICA showed partial overlap (**Table 2**). The overlap was not total in any of the cases, with some pairs showing more correspondence than others. The LC pupil mask showed little overlap with the masks from the structural methods (AT and NM) but some overlap with the mask from the functional method (mICA).

**Table 2.** The overlap was calculated between the atlas mask, the pupil mask, and the group-average NM and mICA masks. The percentage in each cell is based on the size of the mask at the header of the raw; for instance, 35% of the total size of LC AT mask overlapped with LC NM mask, while 38% of the total size of LC NM mask overlapped with LC AT.

| Overlap (*bilateral*) | LC AT | LC NM | LC mICA | LC pupil |
| --- | --- | --- | --- | --- |
| **LC AT** | - | 35% | 23% | 1% |
| **LC NM** | 38% | - | 41% | 1% |
| **LC mICA** | 25% | 40% | - | 23% |
| **LC_pupil** | 1% | 11% | 12% | - |

The peak coordinates of the different masks presented some variability, as indicated by the Euclidean distance between coordinates (**Table 3**). The mICA and pupil masks seemed to reflect a more caudal portion of the LC, since their coordinates are closer to AT2.

**Table 3.** Euclidean distance (mm) between centers of coordinates of the different group masks.

| LC LEFT | AT1 | AT2 | NM | MICA |
| --- | --- | --- | --- | --- |
| **AT1** |  |  |  |  |
| **AT2** | 3.16 |  |  |  |
| **NM** | 2.25 | 1.57 |  |  |
| **MICA** | 3.91 | 1.64 | 2.23 |  |
| **PUPIL** | 6.72 | 3.83 | 4.58 | 3.16 |
| LC RIGHT | AT1 | AT2 | NM | mICA |
| **AT1** |  |  |  |  |
| **AT2** | 3.05 |  |  |  |
| **NM** | 2.37 | 1.42 |  |  |
| **MICA** | 5.23 | 2.87 | 3.6 |  |

We then calculated the coordinates with the most probability of getting a true LC signal. We overlapped the four masks (AT, NM, mICA and pupil; **Figure 5** and **Figure 7**) and retained the voxels included in at least three of the masks.

**Figure 7.**
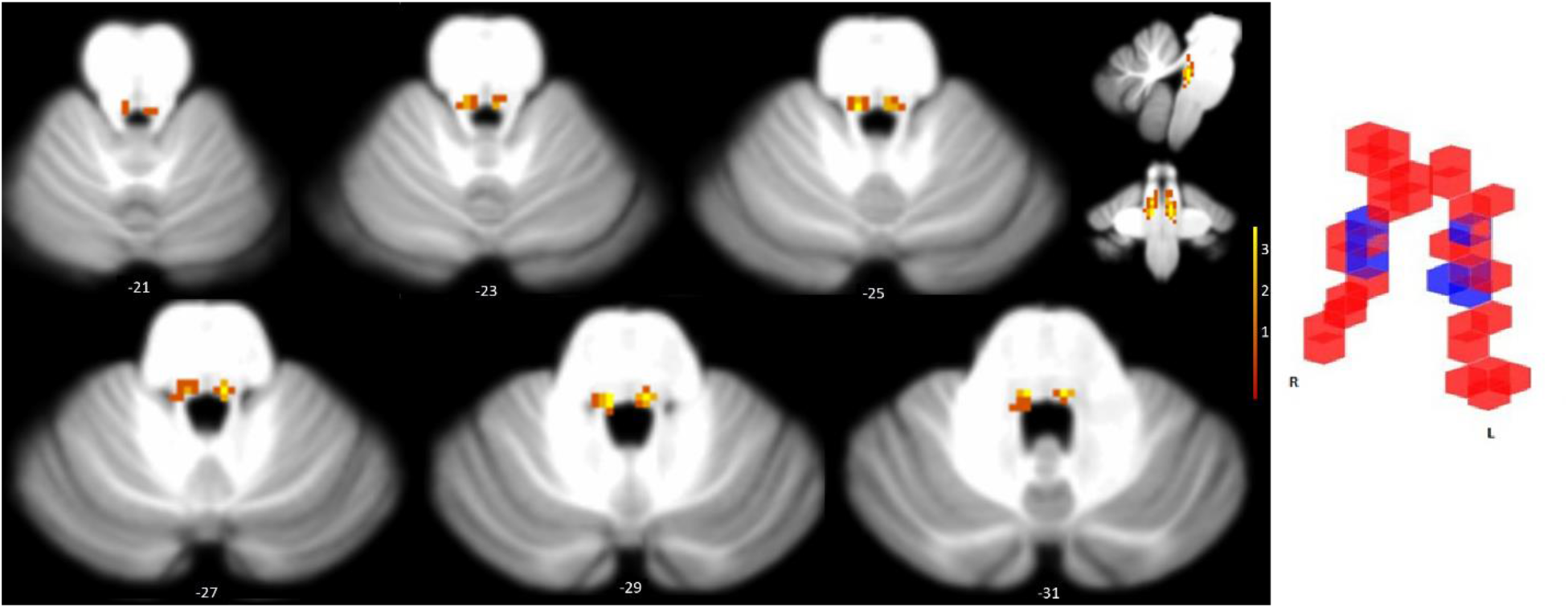
Consensus of LC location at the group level. Left panel: overlap between LC AT, NM, mICA and pupil masks. Left hemisphere on the left. Right panel: voxels showing overlap of at least 3 masks (blue) superimposed on the LC AT mask (red).

### 3.4 LC individual masks

Individual masks presented peak values at locations that more or less closely match the spatial distribution of the NM-based masks (AT and mean NM), but a certain degree of variability can be observed (**Figure 8**). It can be also observed that the mICA individual masks present larger variability than the NM masks.

**Figure 8.**
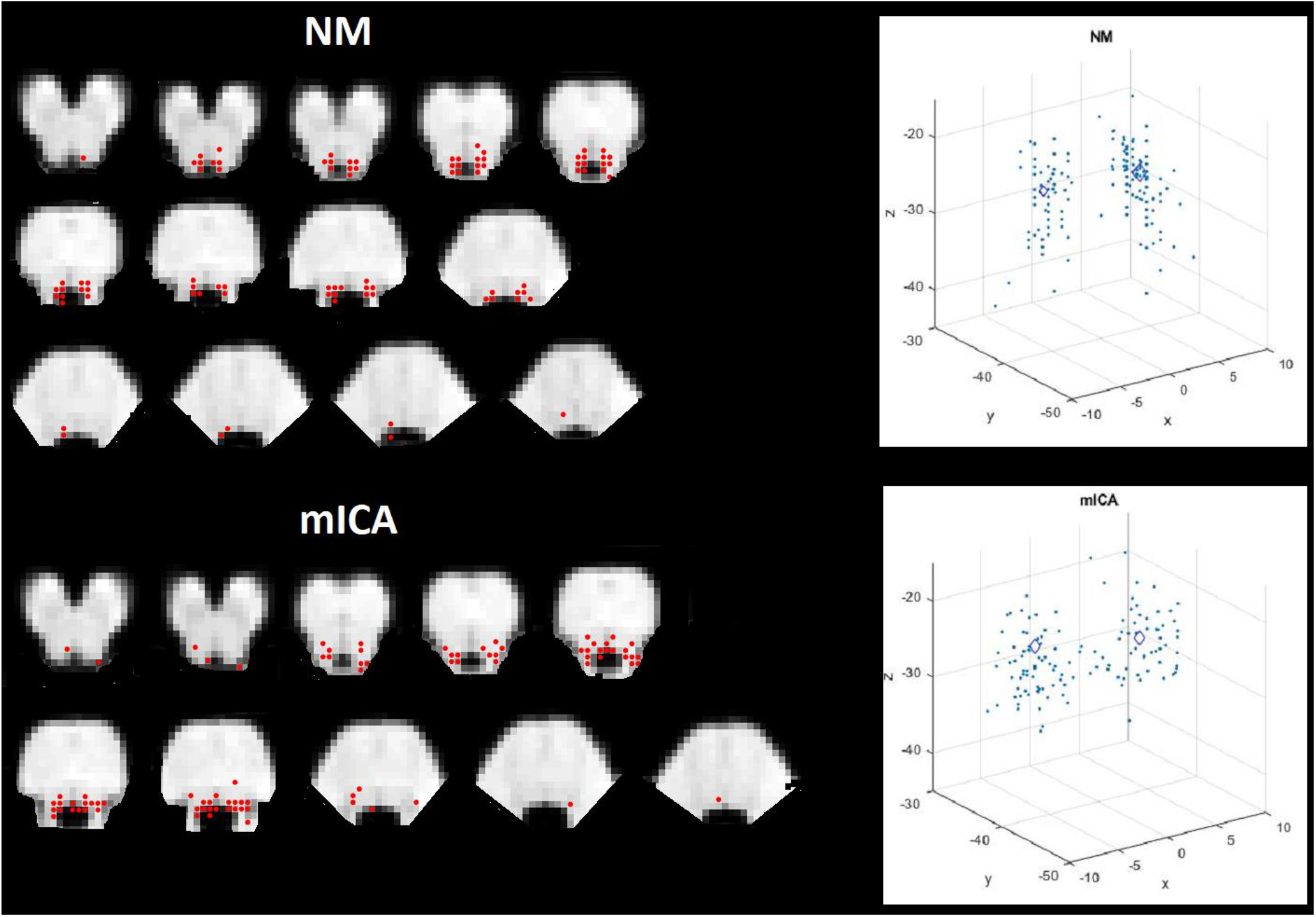
Distribution of coordinates of center of mass of individual LC masks. Top: NM; bottom: mICA. Slices: data overlapped on brainstem template. Scatter plots: data displayed on MNI axis of coordinates. Diamonds indicate mean coordinate, and size of diamonds correspond to dispersion in z axis.

We correlated the signal extracted from the different masks in each individual’s resting state scan. **Table 4** shows the average correlation. We observed an overall positive correlation between LC masks, with values ranging from 0.5 to 0.94. As a confirmation of the validity of our neuromelanin-based approach, the highest correlation was observed between LC AT and mean LC NM, as was expected because they were obtained using equivalent approaches. We observed lowest overall correlation between the pupil mask and the individual masks (NM and mICA), moderate correlation between the individual masks (NM and mICA), and relatively high correlation between the group-average masks (AT, mean NM and consensus) between each other and with the other masks.

**Table 4.**
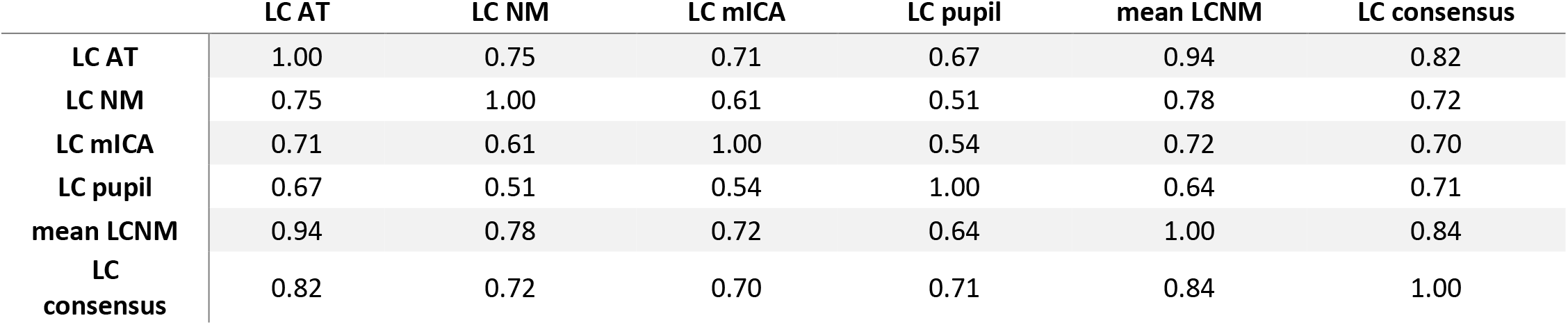
Correlation of signal extracted from the different LC masks.

|  | LC AT | LC NM | LC mICA | LC pupil | mean LCNM | LC consensus |
| --- | --- | --- | --- | --- | --- | --- |
| LC AT | 1.00 | 0.75 | 0.71 | 0.67 | 0.94 | 0.82 |
| LC NM | 0.75 | 1.00 | 0.61 | 0.51 | 0.78 | 0.72 |
| LC mICA | 0.71 | 0.61 | 1.00 | 0.54 | 0.72 | 0.70 |
| LC pupil | 0.67 | 0.51 | 0.54 | 1.00 | 0.64 | 0.71 |
| mean LCNM | 0.94 | 0.78 | 0.72 | 0.64 | 1.00 | 0.84 |
| LC consensus | 0.82 | 0.72 | 0.70 | 0.71 | 0.84 | 1.00 |

### 3.5 Functional connectivity with the different LC seeds

We then obtained the FC patterns associated with each LC seed (**Figure 2** and **Figure 9**). We observed an overall qualitatively similar pattern of connectivity across seeds, with significant clusters in the cerebellum, inferior frontal, parietal and occipital clusters, as well as insula and anterior cingulate. Other regions appeared in some but not all the functional connectivity maps (**Table 5**).

**Figure 9.**
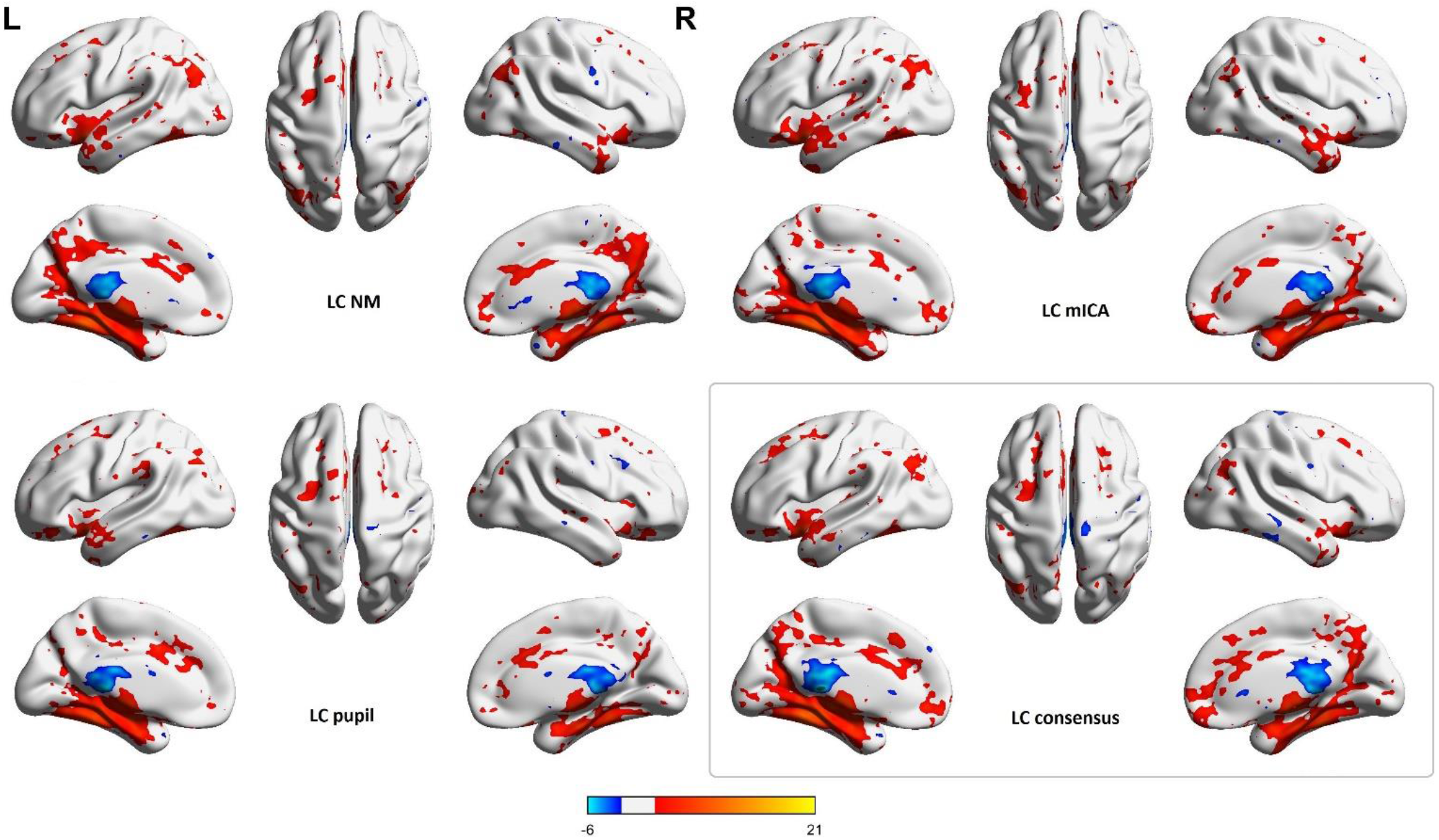
Functional connectivity maps corresponding to the different LC seeds.

**Table 5.**
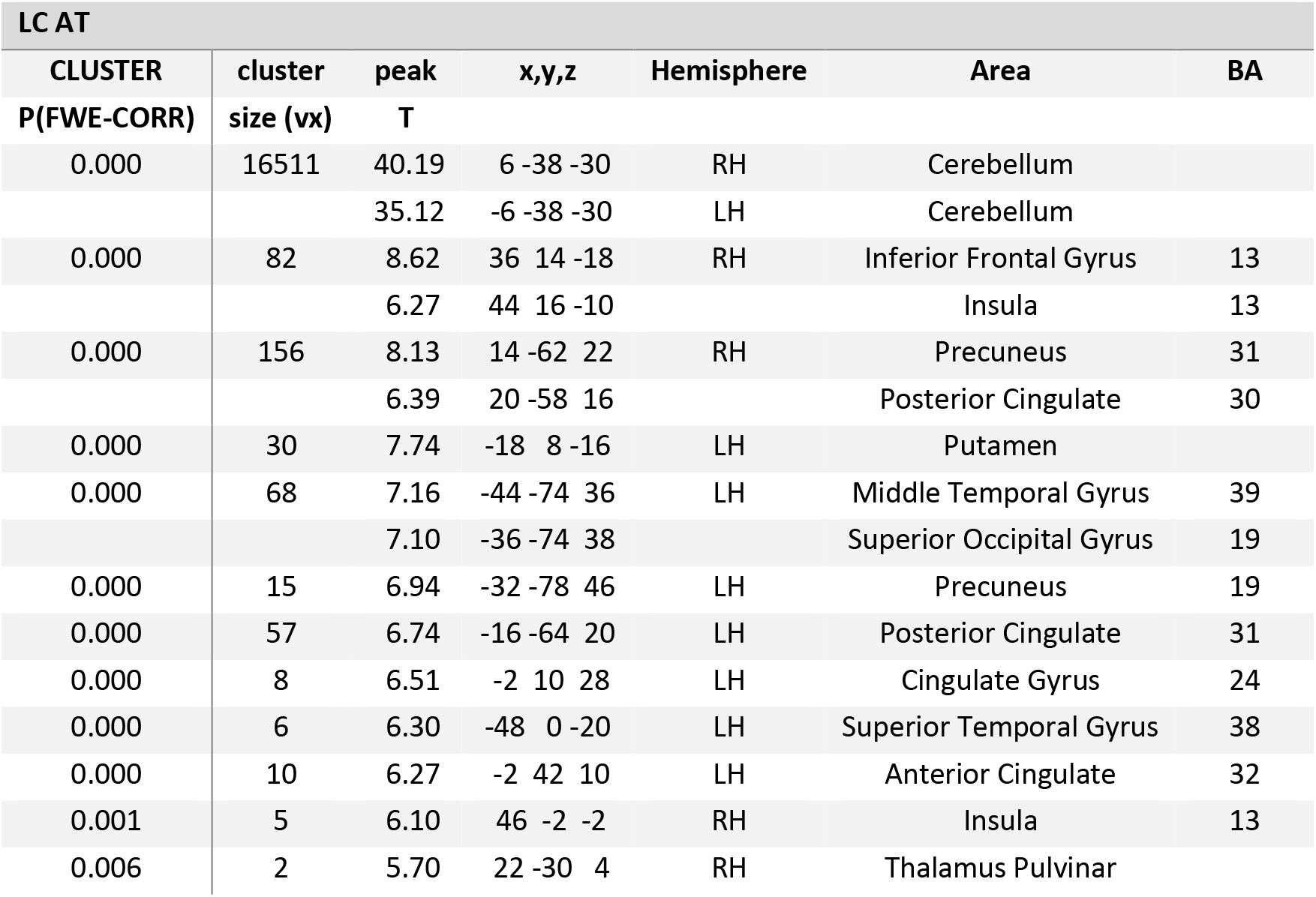
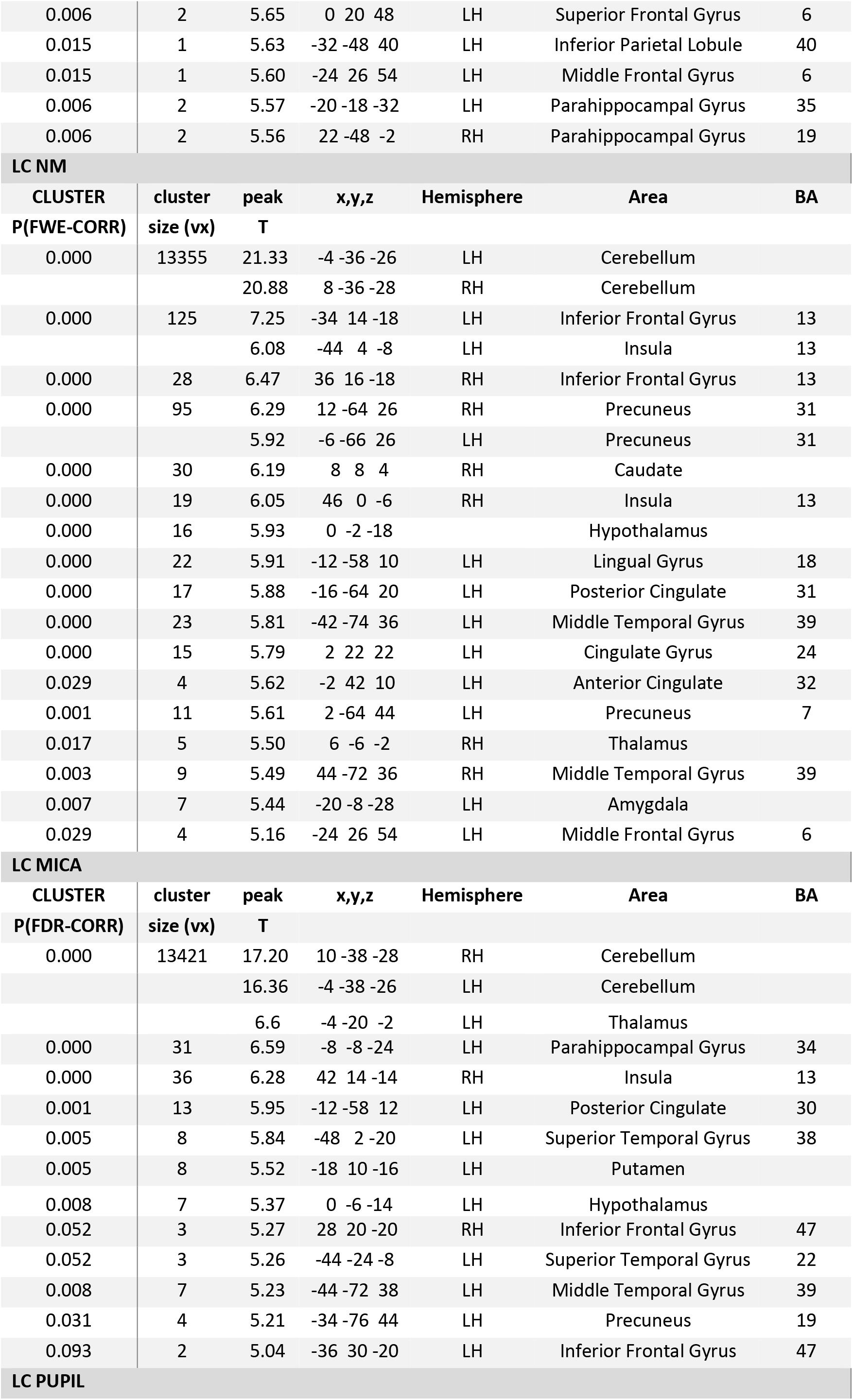
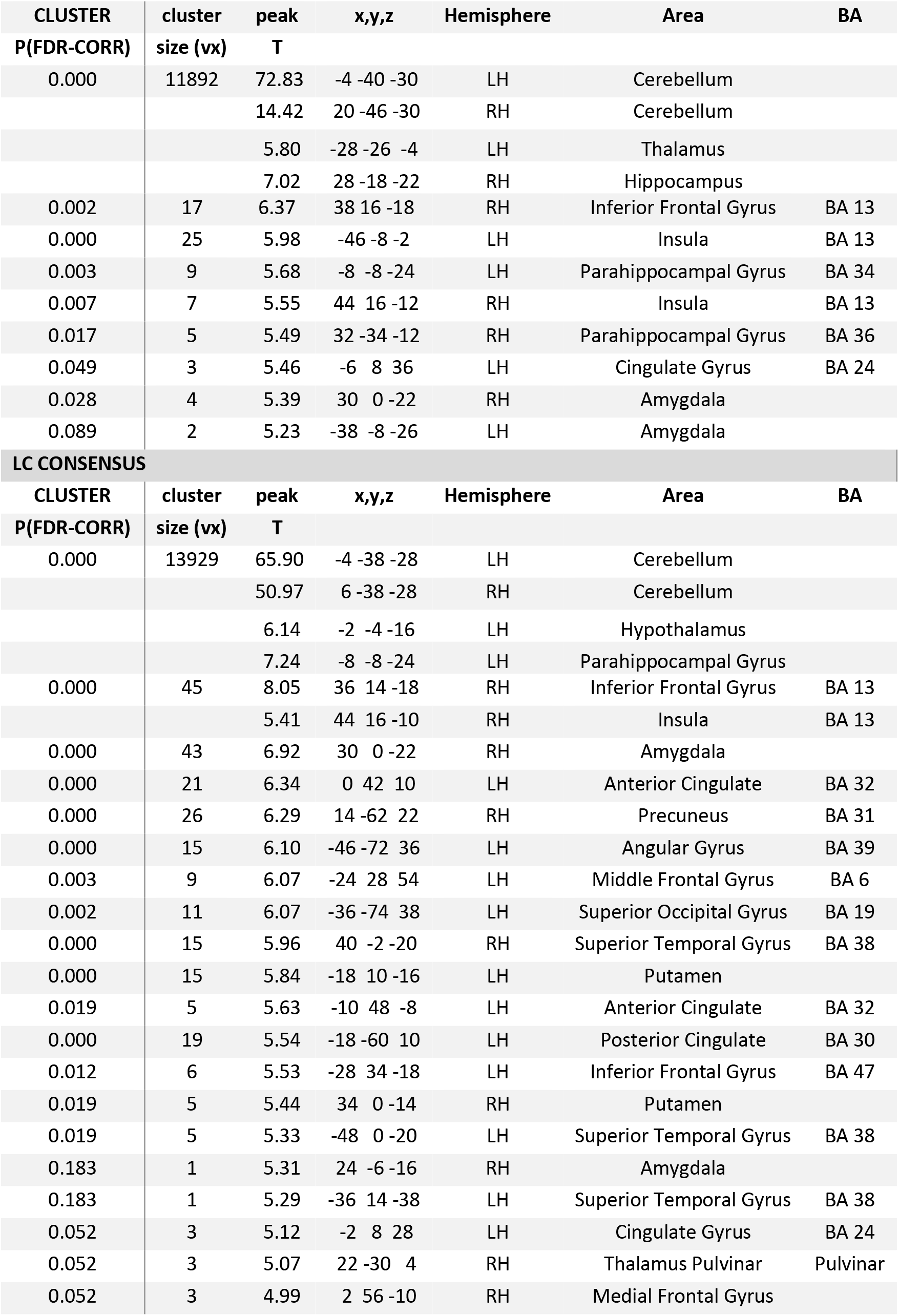
Clusters of significant FC with the LC seeds. LC AT: cluster p < 0.05 (FWE). For the rest, p < 0.5 (FWE)

**Figure 9. Functional connectivity maps corresponding to the different LC seeds.**
| LC AT |  |  |  |  |  |  |
| --- | --- | --- | --- | --- | --- | --- |
| CLUSTER | cluster | peak | x,y,z | Hemisphere | Area | BA |
| P(FWE-CORR) | size (vx) | T |  |  |  |  |
| 0.000 | 16511 | 40.19 | 6 -38 -30 | RH | Cerebellum |  |
|  |  | 35.12 | -6 -38 -30 | LH | Cerebellum |  |
| 0.000 | 82 | 8.62 | 36 14 -18 | RH | Inferior Frontal Gyrus | 13 |
|  |  | 6.27 | 44 16 -10 |  | Insula | 13 |
| 0.000 | 156 | 8.13 | 14 -62 22 | RH | Precuneus | 31 |
|  |  | 6.39 | 20 -58 16 |  | Posterior Cingulate | 30 |
| 0.000 | 30 | 7.74 | -18 8 -16 | LH | Putamen |  |
| 0.000 | 68 | 7.16 | -44 -74 36 | LH | Middle Temporal Gyrus | 39 |
|  |  | 7.10 | -36 -74 38 |  | Superior Occipital Gyrus | 19 |
| 0.000 | 15 | 6.94 | -32 -78 46 | LH | Precuneus | 19 |
| 0.000 | 57 | 6.74 | -16 -64 20 | LH | Posterior Cingulate | 31 |
| 0.000 | 8 | 6.51 | -2 10 28 | LH | Cingulate Gyrus | 24 |
| 0.000 | 6 | 6.30 | -48 0 -20 | LH | Superior Temporal Gyrus | 38 |
| 0.000 | 10 | 6.27 | -2 42 10 | LH | Anterior Cingulate | 32 |
| 0.001 | 5 | 6.10 | 46 -2 -2 | RH | Insula | 13 |
| 0.006 | 2 | 5.70 | 22 -30 4 | RH | Thalamus Pulvinar |  |
| 0.006 | 2 | 5.65 | 0 20 48 | LH | Superior Frontal Gyrus | 6 |
| 0.015 | 1 | 5.63 | -32 -48 40 | LH | Inferior Parietal Lobule | 40 |
| 0.015 | 1 | 5.60 | -24 26 54 | LH | Middle Frontal Gyrus | 6 |
| 0.006 | 2 | 5.57 | -20 -18 -32 | LH | Parahippocampal Gyrus | 35 |
| 0.006 | 2 | 5.56 | 22 -48 -2 | RH | Parahippocampal Gyrus | 19 |

**LC NM**
| **CLUSTER** | **cluster** | **peak** | **x,y,z** | **Hemisphere** | **Area** | **BA** |
| --- | --- | --- | --- | --- | --- | --- |
| **P(FWE-CORR)** | **size (vx)** | **T** |  |  |  |  |
| 0.000 | 13355 | 21.33 | -4 -36 -26 | LH | Cerebellum |  |
|  |  | 20.88 | 8 -36 -28 | RH | Cerebellum |  |
| 0.000 | 125 | 7.25 | -34 14 -18 | LH | Inferior Frontal Gyrus | 13 |
|  |  | 6.08 | -44 4 -8 | LH | Insula | 13 |
| 0.000 | 28 | 6.47 | 36 16 -18 | RH | Inferior Frontal Gyrus | 13 |
| 0.000 | 95 | 6.29 | 12 -64 26 | RH | Precuneus | 31 |
|  |  | 5.92 | -6 -66 26 | LH | Precuneus | 31 |
| 0.000 | 30 | 6.19 | 8 8 4 | RH | Caudate |  |
| 0.000 | 19 | 6.05 | 46 0 -6 | RH | Insula | 13 |
| 0.000 | 16 | 5.93 | 0 -2 -18 |  | Hypothalamus |  |
| 0.000 | 22 | 5.91 | -12 -58 10 | LH | Lingual Gyrus | 18 |
| 0.000 | 17 | 5.88 | -16 -64 20 | LH | Posterior Cingulate | 31 |
| 0.000 | 23 | 5.81 | -42 -74 36 | LH | Middle Temporal Gyrus | 39 |
| 0.000 | 15 | 5.79 | 2 22 22 | LH | Cingulate Gyrus | 24 |
| 0.029 | 4 | 5.62 | -2 42 10 | LH | Anterior Cingulate | 32 |
| 0.001 | 11 | 5.61 | 2 -64 44 | LH | Precuneus | 7 |
| 0.017 | 5 | 5.50 | 6 -6 -2 | RH | Thalamus |  |
| 0.003 | 9 | 5.49 | 44 -72 36 | RH | Middle Temporal Gyrus | 39 |
| 0.007 | 7 | 5.44 | -20 -8 -28 | LH | Amygdala |  |
| 0.029 | 4 | 5.16 | -24 26 54 | LH | Middle Frontal Gyrus | 6 |

**LC MICA**
| **CLUSTER** | **cluster** | **peak** | **x,y,z** | **Hemisphere** | **Area** | **BA** |
| --- | --- | --- | --- | --- | --- | --- |
| **P(FDR-CORR)** | **size (vx)** | **T** |  |  |  |  |
| 0.000 | 13421 | 17.20 | 10 -38 -28 | RH | Cerebellum |  |
|  |  | 16.36 | -4 -38 -26 | LH | Cerebellum |  |
|  |  | 6.6 | -4 -20 -2 | LH | Thalamus |  |
| 0.000 | 31 | 6.59 | -8 -8 -24 | LH | Parahippocampal Gyrus | 34 |
| 0.000 | 36 | 6.28 | 42 14 -14 | RH | Insula | 13 |
| 0.001 | 13 | 5.95 | -12 -58 12 | LH | Posterior Cingulate | 30 |
| 0.005 | 8 | 5.84 | -48 2 -20 | LH | Superior Temporal Gyrus | 38 |
| 0.005 | 8 | 5.52 | -18 10 -16 | LH | Putamen |  |
| 0.008 | 7 | 5.37 | 0 -6 -14 | LH | Hypothalamus |  |
| 0.052 | 3 | 5.27 | 28 20 -20 | RH | Inferior Frontal Gyrus | 47 |
| 0.052 | 3 | 5.26 | -44 -24 -8 | LH | Superior Temporal Gyrus | 22 |
| 0.008 | 7 | 5.23 | -44 -72 38 | LH | Middle Temporal Gyrus | 39 |
| 0.031 | 4 | 5.21 | -34 -76 44 | LH | Precuneus | 19 |
| 0.093 | 2 | 5.04 | -36 30 -20 | LH | Inferior Frontal Gyrus | 47 |

| CLUSTER | cluster | peak | x,y,z | Hemisphere | Area | BA |
| --- | --- | --- | --- | --- | --- | --- |
| P(FDR-CORR) | size (vx) | T |  |  |  |  |
| 0.000 | 11892 | 72.83 | -4 -40 -30 | LH | Cerebellum |  |
|  |  | 14.42 | 20 -46 -30 | RH | Cerebellum |  |
|  |  | 5.80 | -28 -26 -4 | LH | Thalamus |  |
|  |  | 7.02 | 28 -18 -22 | RH | Hippocampus |  |
| 0.002 | 17 | 6.37 | 38 16 -18 | RH | Inferior Frontal Gyrus | BA 13 |
| 0.000 | 25 | 5.98 | -46 -8 -2 | LH | Insula | BA 13 |
| 0.003 | 9 | 5.68 | -8 -8 -24 | LH | Parahippocampal Gyrus | BA 34 |
| 0.007 | 7 | 5.55 | 44 16 -12 | RH | Insula | BA 13 |
| 0.017 | 5 | 5.49 | 32 -34 -12 | RH | Parahippocampal Gyrus | BA 36 |
| 0.049 | 3 | 5.46 | -6 8 36 | LH | Cingulate Gyrus | BA 24 |
| 0.028 | 4 | 5.39 | 30 0 -22 | RH | Amygdala |  |
| 0.089 | 2 | 5.23 | -38 -8 -26 | LH | Amygdala |  |
| **LC CONSENSUS** |  |  |  |  |  |  |
| CLUSTER | cluster | peak | x,y,z | Hemisphere | Area | BA |
| P(FDR-CORR) | size (vx) | T |  |  |  |  |
| 0.000 | 13929 | 65.90 | -4 -38 -28 | LH | Cerebellum |  |
|  |  | 50.97 | 6 -38 -28 | RH | Cerebellum |  |
|  |  | 6.14 | -2 -4 -16 | LH | Hypothalamus |  |
|  |  | 7.24 | -8 -8 -24 | LH | Parahippocampal Gyrus |  |
| 0.000 | 45 | 8.05 | 36 14 -18 | RH | Inferior Frontal Gyrus | BA 13 |
|  |  | 5.41 | 44 16 -10 | RH | Insula | BA 13 |
| 0.000 | 43 | 6.92 | 30 0 -22 | RH | Amygdala |  |
| 0.000 | 21 | 6.34 | 0 42 10 | LH | Anterior Cingulate | BA 32 |
| 0.000 | 26 | 6.29 | 14 -62 22 | RH | Precuneus | BA 31 |
| 0.000 | 15 | 6.10 | -46 -72 36 | LH | Angular Gyrus | BA 39 |
| 0.003 | 9 | 6.07 | -24 28 54 | LH | Middle Frontal Gyrus | BA 6 |
| 0.002 | 11 | 6.07 | -36 -74 38 | LH | Superior Occipital Gyrus | BA 19 |
| 0.000 | 15 | 5.96 | 40 -2 -20 | RH | Superior Temporal Gyrus | BA 38 |
| 0.000 | 15 | 5.84 | -18 10 -16 | LH | Putamen |  |
| 0.019 | 5 | 5.63 | -10 48 -8 | LH | Anterior Cingulate | BA 32 |
| 0.000 | 19 | 5.54 | -18 -60 10 | LH | Posterior Cingulate | BA 30 |
| 0.012 | 6 | 5.53 | -28 34 -18 | LH | Inferior Frontal Gyrus | BA 47 |
| 0.019 | 5 | 5.44 | 34 0 -14 | RH | Putamen |  |
| 0.019 | 5 | 5.33 | -48 0 -20 | LH | Superior Temporal Gyrus | BA 38 |
| 0.183 | 1 | 5.31 | 24 -6 -16 | RH | Amygdala |  |
| 0.183 | 1 | 5.29 | -36 14 -38 | LH | Superior Temporal Gyrus | BA 38 |
| 0.052 | 3 | 5.12 | -2 8 28 | LH | Cingulate Gyrus | BA 24 |
| 0.052 | 3 | 5.07 | 22 -30 4 | RH | Thalamus Pulvinar | Pulvinar |
| 0.052 | 3 | 4.99 | 2 56 -10 | RH | Medial Frontal Gyrus |  |

To examine the similarity between connectivity maps from the different seeds, we calculated the Pearson correlation between the maps of each individual (**Figure 10**, partial correlation with respect to the FC map with PT as seed). Average correlation values ranged from 0.33 (LC AT-pupil) to 0.93 (LC AT-mean NM). The highest similarity was obtained between the FC maps from LC AT and mean LC NM; which was expected because both masks were obtained using the same approach. High congruence was observed between the maps from LC AT and the maps from NM and mICA (not significantly different from each other after correcting for multiple comparisons, p = 0.008). The maps from LC mICA and LC NM showed the next highest similarity, followed by the maps from LC pupil and the other study-specific masks. The fact that the similarity between the maps of LC pupil with LC NM and with mICA (not significantly different from each other, p = 0.164) were higher as compared to LC AT (pupil-NM and pupil-mICA vs pupil-AT: p < 10^−10^) may indicate that it is capturing an LC component of this particular sample that LC AT is not capturing. Lastly, the FC map from the consensus mask showed high similarity with those of all the other masks (**Figure 10**). The similarity between mean LC consensus and LC NM was highest (p < 10^−5^), followed by LC AT (p < 10^−7^). The similarity between mean LC consensus and the other masks was not different from each other (p > 0.2).

**Figure 10.**
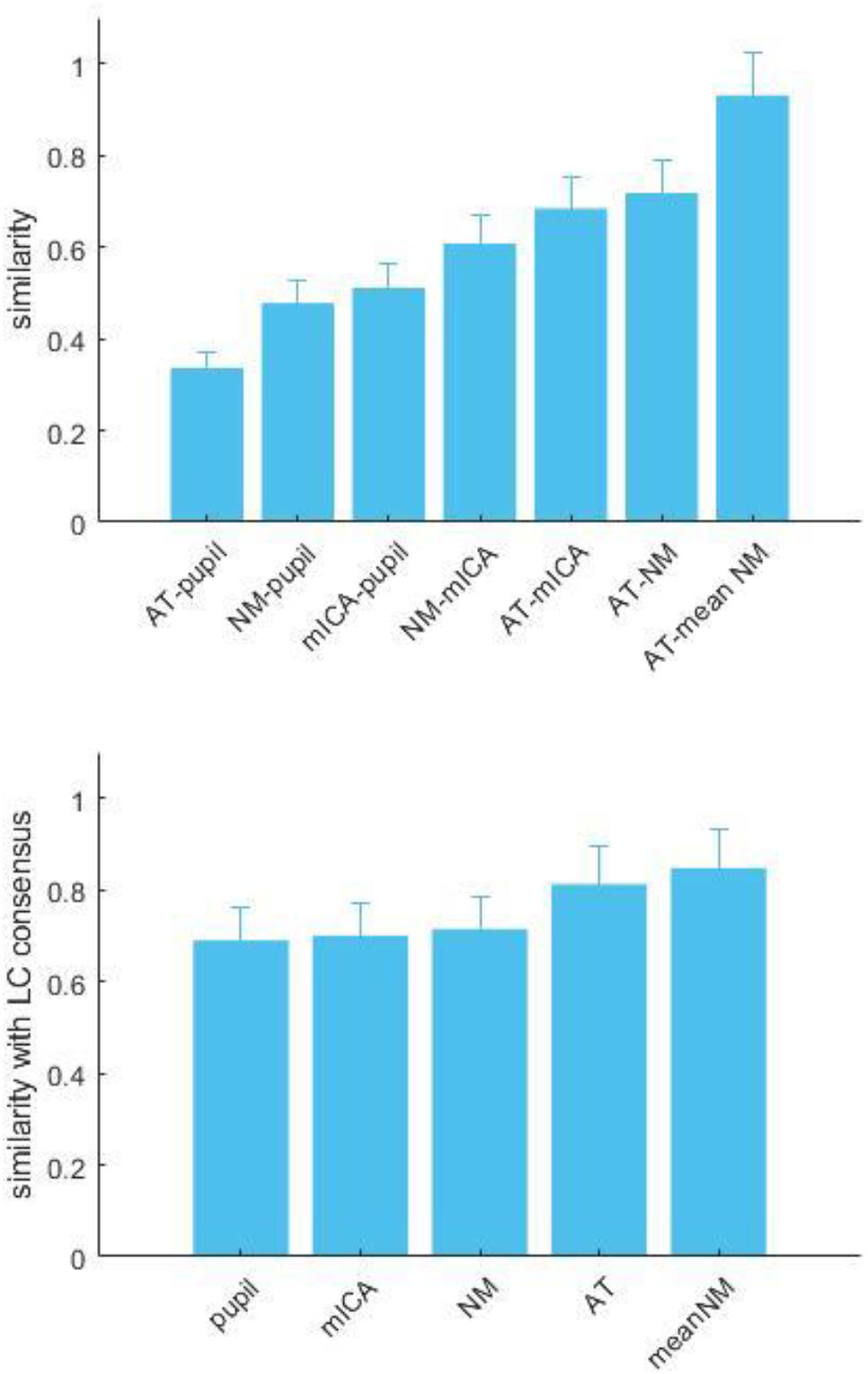
Similarity between the FC maps from the different seeds. Top: similarity between maps of different seeds, as indicated by the x labels. Bottom: similarity of the FC maps from the different seeds with the FC map obtained from the LC consensus mask.

To further characterize the similarity between FC maps between seeds at the individual level, we did two further analysis. We created Bland-Altman plots to compare the similarity of the *statistical values* of the maps. The plots showed that, across thresholds, there was a tendency for the AT mask to depart towards larger T values (indicated by the difference in the y axis), compared to NM, mICA and pupil (**Figure 11**). They also showed an agreement between the NM and mICA maps, as shown by points lying around the zero of the y axis. Pupil maps also showed differences at larger T values with respect to NM and mICA maps.

**Figure 11.**
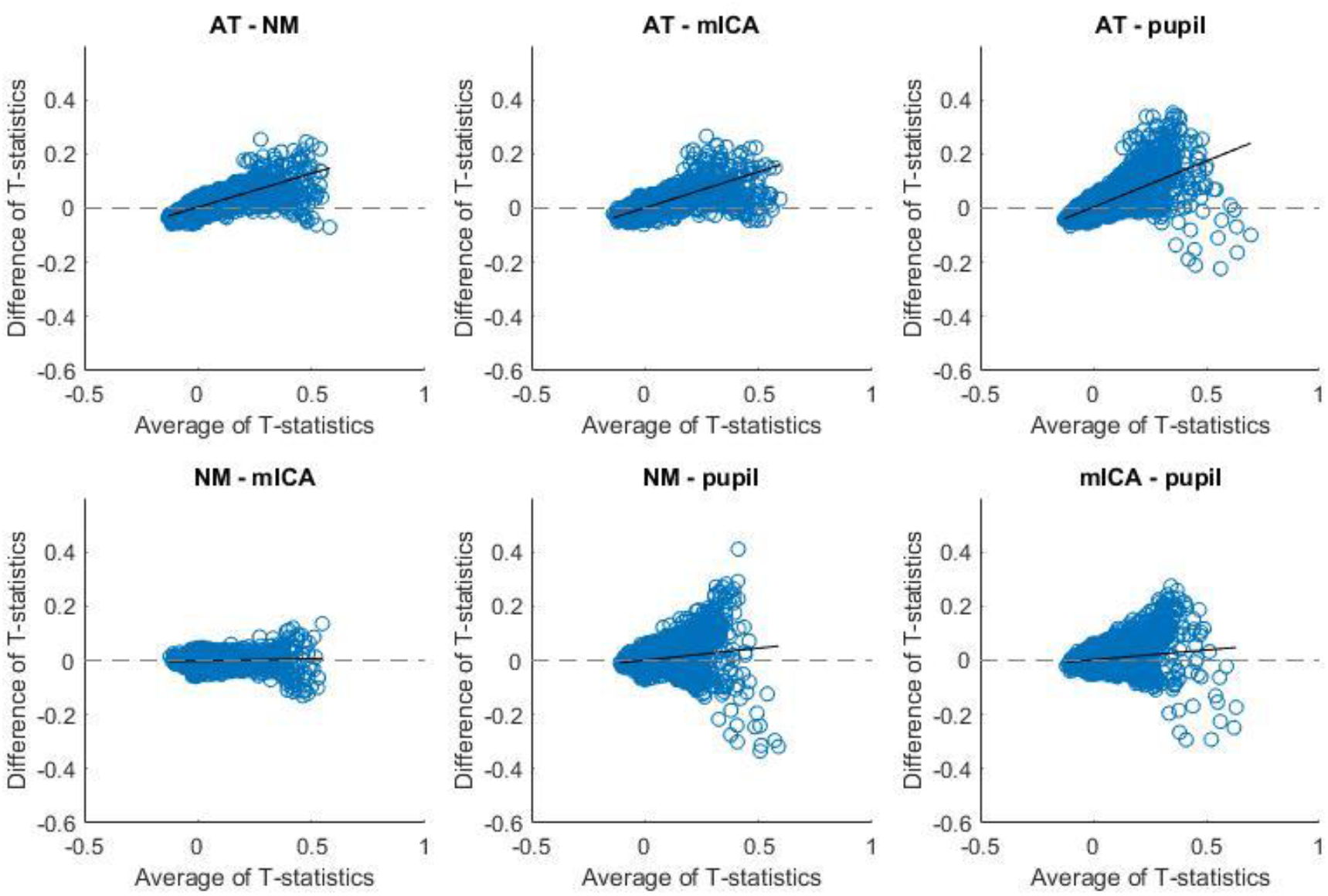
Bland-Altman plots.

We then calculated the Euler characteristic (EC) of the maps at different thresholds. The EC and cluster count measures (**Figure 12**) give an idea of the number of clusters and its relation with the “holes” between them in the maps at the different T thresholds. Therefore, this method is sensitive to an overall concordance in spatial distribution despite differences in scale between images. Of interest in these plots is where the peak occurs for each statistical map. The most evident difference is that the LC AT mask, different from the other three masks, shows a more delayed peak at larger statistical values, indicating differences in topology between them. The cluster count plot indicates that LC AT contains larger clusters (because of the smaller cluster count at lower T) and higher statistical values (because of the slower descend at higher T values) than the other masks.

**Figure 12.**
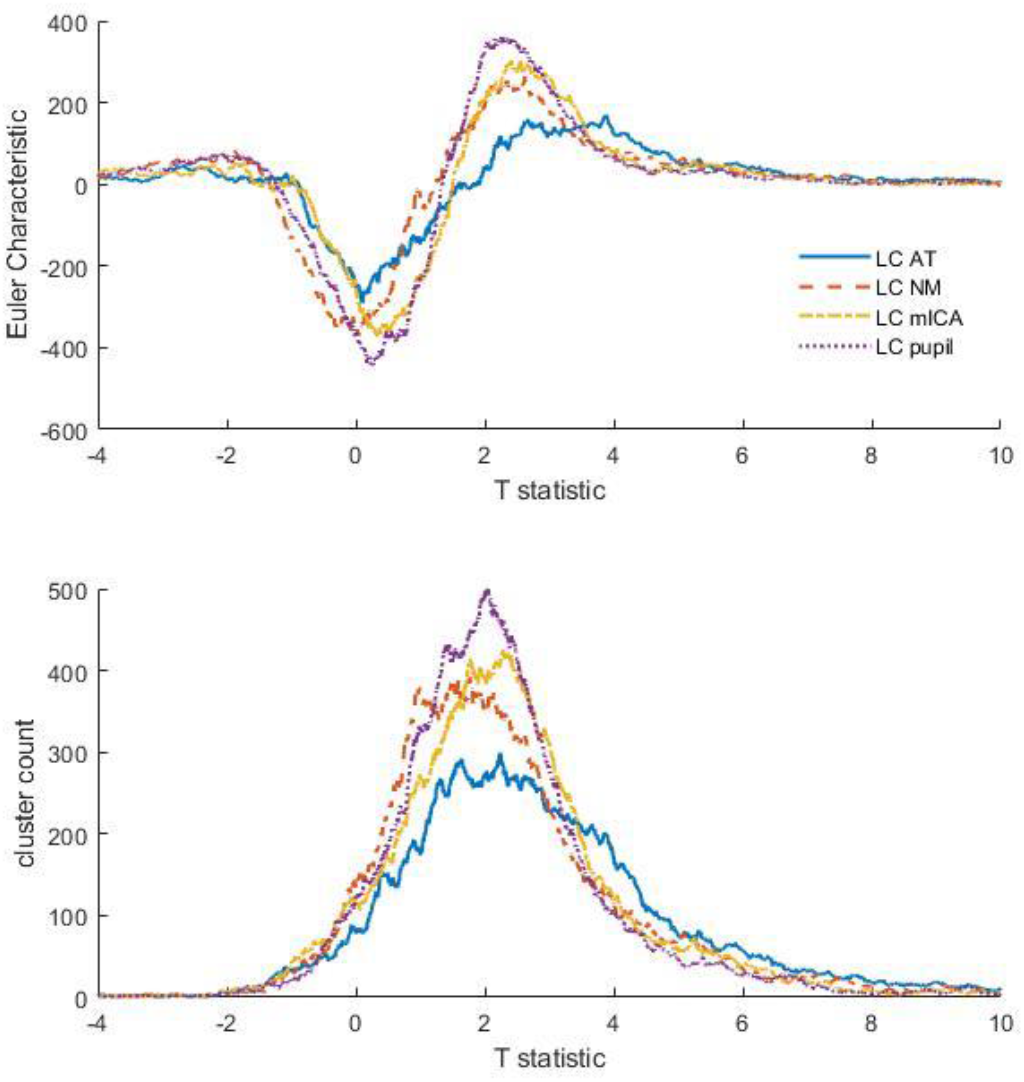
Euler characteristic (top) and cluster count (bottom).

## 4. Discussion

In the current study, we compared different methods of defining locus coeruleus localization for functional analysis. The methods were: a probabilistic map from a previous study (LC AT), manual delineation in neuromelanin-based structural sequence (LC NM), or semi-automatic segmentation in the same sequence (Supplementary material), an independent-components analysis-based signal extraction at the individual level (LC mICA), and regressed pupil activity (LC pupil). We obtained the LC seeds based on the different methods in a large sample, and performed connectivity analysis on resting state data of the same sample. Our main goal was to work towards an assessment of the fidelity of the different LC localization methods, given the large variability in approaches existing in the literature. In our sample, we compared quantitatively the above-mentioned methods as valid localizers of LC, assessed the reliability of connectivity patterns associated to the nucleus’ activity, and examined pipelines for analysis. In **Table 6**, we present different methodological aspects of each approach. The results constitute an advancement for the study of LC activity in humans to further understand its role in health and disease.

**Table 6.**

| METHOD | TIME INVESTED<br>(INCLUDING AMOUNT OF<br>MANUAL WORK) | ADVANTAGES | DISADVANTAGES |
| --- | --- | --- | --- |
| LC AT | + | Good for big datasets. | No individual specificity. |
| LC NM | +++ | Allows individual localization. Good for small datasets.<br>Allows extraction of volumetric information /signal intensity. | Additional scanning time. Manual procedure. May suffer changes with age. |
| LC NM SS | ++ | Same as above. | Additional scanning time. May suffer changes with age. |
| LC MICA | ++++ | Allows individual localization when a NM scan is not available. Easier to detect lateral LC. | Includes manual steps. |
| LC PUPIL | ++ | Functionally defined LC localization. | Poor individual specificity. |

### 4.1 Group-level LC masks

We identified two main applications for LC localization with functional data (Figure 1). One was to get a group-level, sample-specific, average mask of LC. This could be necessary for, for example, checking LC activation on group statistical maps (second-level analysis). The other was to obtain individual LC masks with the highest spatial specificity possible. This could be used, for example, to extract BOLD activity. For the first approach, we obtained an average of individual NM masks, a mask based on group-level significant pupil signal regression, and a consensus mask across these LC masks. This approach is the most conservative. For the second approach, we compared the functional similarity (correlation of activity and resulting functional connectivity maps) of the individual masks obtained from the different approaches.

The coordinates from the different average masks fell close to the medium portion of LC (Figure 6). This portion shows the least variability across individuals and the largest voxel count after threshold-based segmentation (Ye et al., 2020). The peaks of the average masks were somehow close to each other (Table 3). Despite this overall concordance, the LC pupil mask was more caudal than the others. Both the LC pupil mask and the average mICA mask were closer to the more caudal peak of the LC AT (z = −27), indicating that these functional-based methods may capture a different functional portion of LC than the structural methods.

Our consensus mask (Figure 7) is within the area of largest overlap between probabilistic maps of previously reported studies, between z = −29 and z = −23 (Dahl et al., 2019; Keren et al., 2009; Tona et al., 2017). Furthermore, the location of our consensus mask agrees with the center of distribution of center coordinates from functional studies, as reported by the meta-analysis of Liu et al. 2017. This suggests that the consensus region may provide a “safe” region as to conclude whether LC was reliably activated in a group statistical map (given that brainstem regions of functional images had good alignment).

### 4.2 LC pupil mask

The presence of a cluster in LC that correlated with pupil size was expected from animal studies and fMRI studies (Alnæs et al., 2014; Joshi et al., 2016; Murphy et al., 2014). The pupillometry-based mask was unilateral at the specified statistical threshold. This is consistent with the results from Murphy et al. 2014, where pupil correlation during rest was found in the left LC at a more caudal portion (z = −30), while during task the peak fell in the right hemisphere at a more rostral portion (z = −22). Schneider et al. 2016 reported a cluster of correlation with pupil change (not pupil size) in the left hemisphere at z = −30 (Table 3 of the paper). The LC pupil mask showed little overlap with the structural masks (AT and NM) and some overlap with LC mICA. Taken together, these results suggest that pupil signal may be related to a functional-specific portion of LC in humans. This possibility is worth exploring in future studies since functional specialization within LC has been suggested by recent animal studies (for a review see Chandler et al. 2019).

### 4.3 Individual-level LC masks

For the second approach, we compared two neuromelanin-based segmentations of LC (manual and semi-automatic), and a brainstem-specific independent components-based identification of LC (mICA). The neuromelanin-based masks had an average size of 25 mm^3^ (unilateral), which is comparable to the volume reported in other studies where LC masks were drawn using these scans (Castellanos et al., 2015; Langley et al., 2017; Tona et al., 2017). These masks are smaller than the expected size of LC, but capture the main concentration of noradrenaline-producing cells (Keren et al., 2015). Our results are in agreement with previously reported results from either segmentation method, for instance peak location, average volume, and supremacy of central part of LC (Betts et al., 2017; Turker et al., 2019; Ye et al., 2020). Semi-automatic segmentation has been used in the literature to identify intensity peaks across slices of NM-sensitive scans (Betts et al., 2017; Dahl et al., 2019). The purpose of these studies was to compare LC integrity across age or neurological conditions. In functional studies, LC segmentation was mainly done through manual drawing (Clewett et al., 2018; Krebs et al., 2018; Mäki-Marttunen et al., 2019b; Mäki-Marttunen et al., 2020). Functional analyses benefit from regions that cover the whole area of interest where BOLD signal is averaged, while structural studies often rely on the identification of the voxels with highest intensity in different slices or even in a single slice. Here we used both approaches to obtain individual regions of interest. Overall, the results from the masks from the manual and from the semi-automatic segmentation were highly concordant. The corresponding group average masks showed more overlap with each other than, for instance, the LC AT mask, and the functional connectivity maps showed high similarity. This suggests that the two methods are comparably reliable in identifying LC and related connectivity patterns.

The mICA analysis was based on the identification of a component which peak was closest to the expected LC localization. The distribution of peak coordinates across individuals showed larger dispersion in the mICA approach than the NM approach (Figure 8), including more coverage of more caudal portions. Furthermore, the left and right LC were often separate components, which may be indicative of a certain functional differentiation.

### 4.4 Functional connectivity patterns

FC maps using the different seeds showed consistent connectivity between LC with a set of regions. The cerebellum showed the strongest connectivity, as is commonly reported (for a review, see Liu et al. 2017). Significant clusters in the insula and anterior cingulate gyrus were found across FC maps, which is consistent with the involvement of LC in the salience detection network, of which these areas are key components (Corbetta et al., 2008). Other cortical areas were inferior or middle frontal gyrus, precuneus, middle temporal gyrus and posterior cingulate. Some subcortical areas were present in some of the FC maps, such as putamen, hippocampus and para-hippocampus. These areas have also been reported in previous studies, but not consistently (Liu et al., 2017). Clusters in the amygdala appeared in maps from either structural (LC NM) and functional (LC pupil) seeds, as well as in the consensus mask. This replicates previous studies using spherical seeds (Bär et al., 2016; Metzger et al., 2016) and is consistent with a functional connection between these two structures (Valentino & Van Bockstaele, 2008; Van Bockstaele et al., 1998).

The individual FC patterns obtained from the different seeds showed variation in similarity (Figure 10). LC AT masks showed the largest similarity with the other maps and LC pupil, lowest. The high similarity between the maps of AT mask and the others may indicate that it successfully captures LC signal from individual LC. However, the extent of the FC maps with LC AT as seed showed more widespread connectivity and the statistical values were larger than the other FC maps (Figure 11 and 12), which could be interpreted as the AT mask being not very specific and capturing signal from surrounding regions, as suggested also by Turker et al. 2019.

We consider that the pattern of similarity between maps from the LC consensus mask and the other maps (Figure 10) strongly indicates that: 1) the consensus mask captured true LC signal that swept the differences inherent to each different approach, indicated by the low variability across similarity values between LC consensus and the other masks; 2) the masks from atlas-based approaches (LC AT and mean LC NM) are able to capture most of true LC signal, as indicated by the highest similarity between the maps of these masks with the map of LC consensus, but also 3) the consensus mask was able to capture sample-specific high-probability LC voxels, indicated by the similarly high correlation between the LC consensus maps and all the other sample specific masks (NM, mICA and pupil), which showed variable correlation with each other.

### 4.5 Functional subdivisions of LC

Changes in LC intensity have been noted with age at specific portions of LC, emphasizing a rostro-caudal axis (Dahl et al., 2019; Ye et al., 2020). Corticotropin-releasing factor (CRF) terminals targeting mostly rostral LC (Van Bockstaele et al., 1996) associate this functional differentiation of LC to the response to stress, and in particular to the shutting down of the stress response. This may have important implications for the study of the conditions arising from uncontrollable stress, such as post-traumatic stress disorder, anxiety or psychosis (Atzori et al., 2016; Mäki-Marttunen et al., 2019a; Winklewski et al., 2017). Animal studies have further unveiled an inhomogeneity in LC organization. The distribution of alpha-1 and alpha-2 receptors seem to follow a rostro-caudal distribution, and projections to thalamus vs. hypothalamus appear to follow a similar axis (reviewed in Schwarz and Luo 2015). Interestingly, we found connectivity with hypothalamus and thalamus in most of the FC maps, but in particular structural seeds (which presented a more rostral peak) showed FC with hypothalamus (e.g. LC SS), while functional seeds (which presented a more caudal peak) showed FC with thalamus (e.g. LC pupil). The fact that functional and structural masks may emphasize different subcomponents of LC may have therefore practical implications for the study of LC functional diversity. Furthermore, the comparison of LC localization approaches on older populations and groups where LC is implicated (such as Parkinson’s disease and Alzehimer’s disease), is an interesting avenue of research.

Some of our findings agree with previous suggestions on structural and functional lateralization of the LC: peak coordinates of largest overlap in the group-specific masks (NM and mICA, Table 1) were in the right hemisphere, while the cluster of regression with the pupil was in the left LC. Functional studies on left vs. right LC could in the future reveal important aspects of LC function.

### 4.6 LC intensity values and correlation with age

We obtained a distribution of intensity values in LC (as quantified from the neuromelanin scans) centered in a CNR value of 0.0991. Previous studies used a variety of measures of contrast in LC (for a review, see Langley et al. 2017). Our result is within a standard deviation range of the CNR values reported for healthy adults (Betts et al., 2017; Clewett et al., 2016; Sasaki et al., 2006; Shibata et al., 2006; Tona et al., 2017). Furthermore, we found no significant relation between CNR and age. The relation between LC intensity and age, although found in several studies, could not be replicated by others (Liu et al., 2019; Shibata et al., 2006). On the other hand, we had a relatively short age range in our sample, which may be a reason why we did not find a significant correlation with age.

### 4.7 Clinical implications

A number of studies have come out recently on how LC is implicated in cognitive deficits due to aging or neurological conditions (Betts et al. 2019, Mather and Harley 2016). These results may bear great potential, for instance if biomarkers of neuropsychiatric conditions can be developed based on imaging measurements. Of key importance, these conditions present large inter-individual variability. An individual-based approach is likely more appropriate for clinical application than a group-based approach, unless homogeneity of the degenerative or pathological changes across patients is demonstrated. While most work has addressed LC integrity, that is, intensity in neuromelanin-sensitive scans, functional studies are also of great interest. Our and other labs’ work suggest that individual-based approaches, although implicating more work load, give more similar within-sample and within-subject functional connectivity maps.

### 4.8 Limitations

We acknowledge a series of limitations. First, LC is a very small nucleus. While this is part of the reason why methods of improved localization are needed (such as those compared in this paper), technical issues inherent to its study must be considered. Structures of small size involve several disadvantages: increased partial volume effects, susceptibility to artifacts, and low signal-to-noise ratio. Here, we intended to minimize these confounds by e.g. having a large sample, following careful coregistration and quality-checking steps, and using a small smoothing Kernel on the functional images. Recent studies have also reported good coregistration with specialized tools, e.g. the ANTS toolbox. Second, the LC pupil mask was a group mask. We examined individual (first level) pupil-based connectivity maps but did not find consistently the presence of clusters in LC. While this would have been a great advantage to compare with the other individual LC masks, other approaches may be needed (e.g. larger number of pupil-fMRI samples, or use of tasks known to boost LC activity and pupil size). Third, although our goal was to compare possible methods to extract activity from LC, we did not include all possible methods as found in bibliography. For instance, as reviewed by Liu et al. 2017, some studies defined spheres or boxes around previously reported coordinates. However, we believed that the methods presented here (LC AT, LC NM, LC mICA, LC pupil) would be much superior in individual localization, and therefore compare between them. Fourth, 7T studies have better resolution (Priovoulos et al., 2018; Tona et al., 2019). However, our study on a 3T scanner is useful because it is the most commonly accessible facility for most research groups. In addition, we replicated the main finding of studies made on 7T scanners and post mortem studies (Fernandes et al., 2012). For instance, we found that the central portion of LC, across methods, presents the largest group overlap, while the rostral and caudal portions present les overlap.

## 5. Conclusions

Based on our results, we conclude that the use of different methods to identify LC for functional analysis are not fully equivalent and therefore interpretation should be done with care. In particular, for the purpose of group-level localization, studies may benefit for a multi-modal approach and the extraction of a group-specific consensus mask. In principle, the use of regions of interest based on coordinates from previous studies may not be as reliable as would be desired for conclusive results. For individual-based approaches, the NM method is probably the best one possible, but it may still be missing portions of LC, for instance more caudal parts that may have functional significance. Improvements in NM imaging (e.g. implementation in high field scanners) may help overcome this limitation. Despite showing more variability, mICA method may be still a valid approach for functional datasets where no NM scan has been acquired. The results presented here offer guidance for future functional studies on the relevance of LC-NE system in human cognition.

## Footnotes

1 From literature, the expected size is between 50 and 75 mm^3^ (unilateral; Keren et al. 2015).

